## Supplementary material for "Control of 3ʹ splice site selection by the yeast splicing factor Fyv6": Figure Supplements and Supplementary Tables

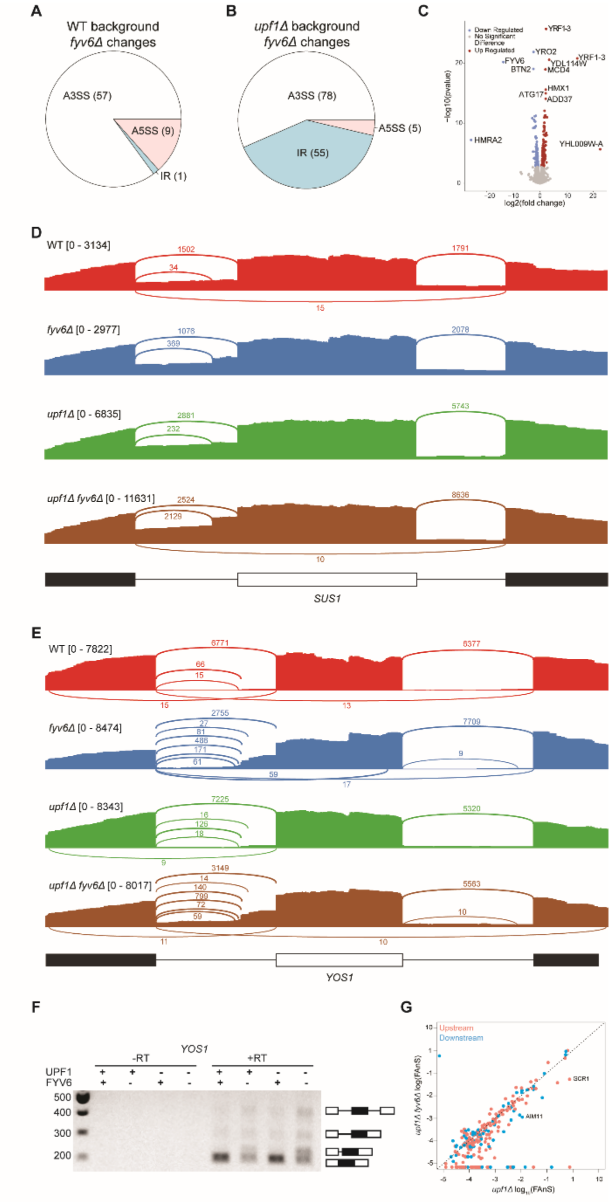


**Figure 1-figure supplement 1. Gene expression analysis based on the RNA-Seq results and an example of Fyv6-dependent splicing changes in *YOS1*. A, B)** Pie charts comparing alternative splicing events discovered by SpliceWiz at 10 PSI in (A) *fyv6Δ* and **(**B) *upf1Δ fyv6Δ* strain backgrounds relative to when Fyv6 is present**.** A3SS = alternate 3' SS; A5SS = alternate 5' SS; IR = intron retention. Note that these data sets were collected at higher read depth than those shown in Fig. 1. As a result, the number of detected changes in splicing cannot be directly compared between the two. **C)** Differential expression analysis in a *upf1Δ* background with presence or absence of Fyv6. Colored points are significant with $p \leq 0.05$ with blue points being downregulated gene expression and red being upregulated. **D)** Sashimi plots showing coverage across splice junctions of *SUS1* in WT (red), *fyv6Δ* (blue), *upf1Δ* (green), and *fyv6Δ upf1Δ* (brown). **E)** Sashimi plots showing coverage across splice junctions of *YOS1* in WT (red), *fyv6Δ* (blue), *upf1Δ* (green), and *fyv6Δ upf1Δ* (brown). **F)** RT-PCR of *YOS1* mRNA showing a higher molecular weight band in strains lacking *FYV6* corresponding to use of an alternative 3' SS in the first intron. **G**) FAnS plot for alternative 5' SS junctions with canonical 3' SS in *upf1Δ* compared to *fyv6Δ upf1Δ.*

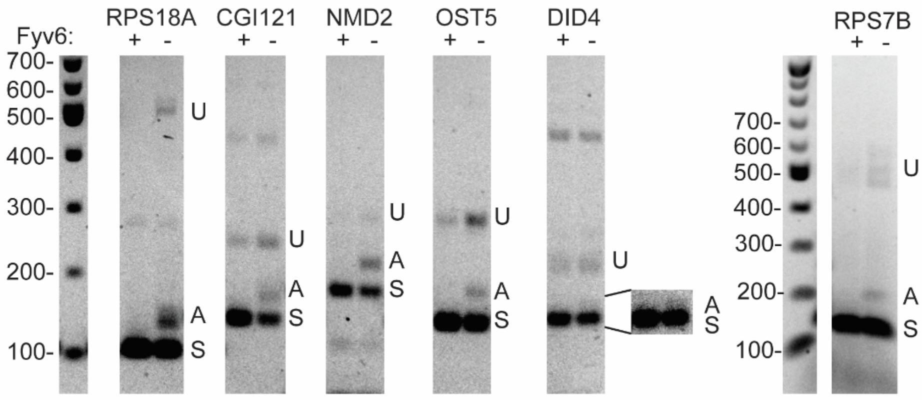


U = unspliced A = Alternative 3′ SS S = spliced

Expected product sizes:

| Gene | Unspliced (bp) | Spliced (bp) | Alternative 3′ SS |
|---|---|---|---|
| RPS18A | 540 | 105 | 135 |
| CGI121 | 251 | 145 | 168 |
| NMD2 | 296 | 183 | 219 |
| OST5 | 293 | 144 | 174 |
| DID4 | 219 | 151 | 158 |
| RPS7B | 491 | 146 | 168 |

**Figure 2-figure supplement 1. Examples of Fyv6-dependent alternative 3′ SS usage.** RT-PCR of RNAs from *upf1Δ* and *fyv6Δ upf1Δ* strains targeting introns in genes identified in the RNA-seq data as having alternative 3′ SS with high FAnS ratios in *fyv6Δ upf1Δ* (see Fig. 2H). The bands for the unspliced, spliced, and alternative 3′ SS products are indicated with U, S, and A, respectively, and their expected sizes are listed.

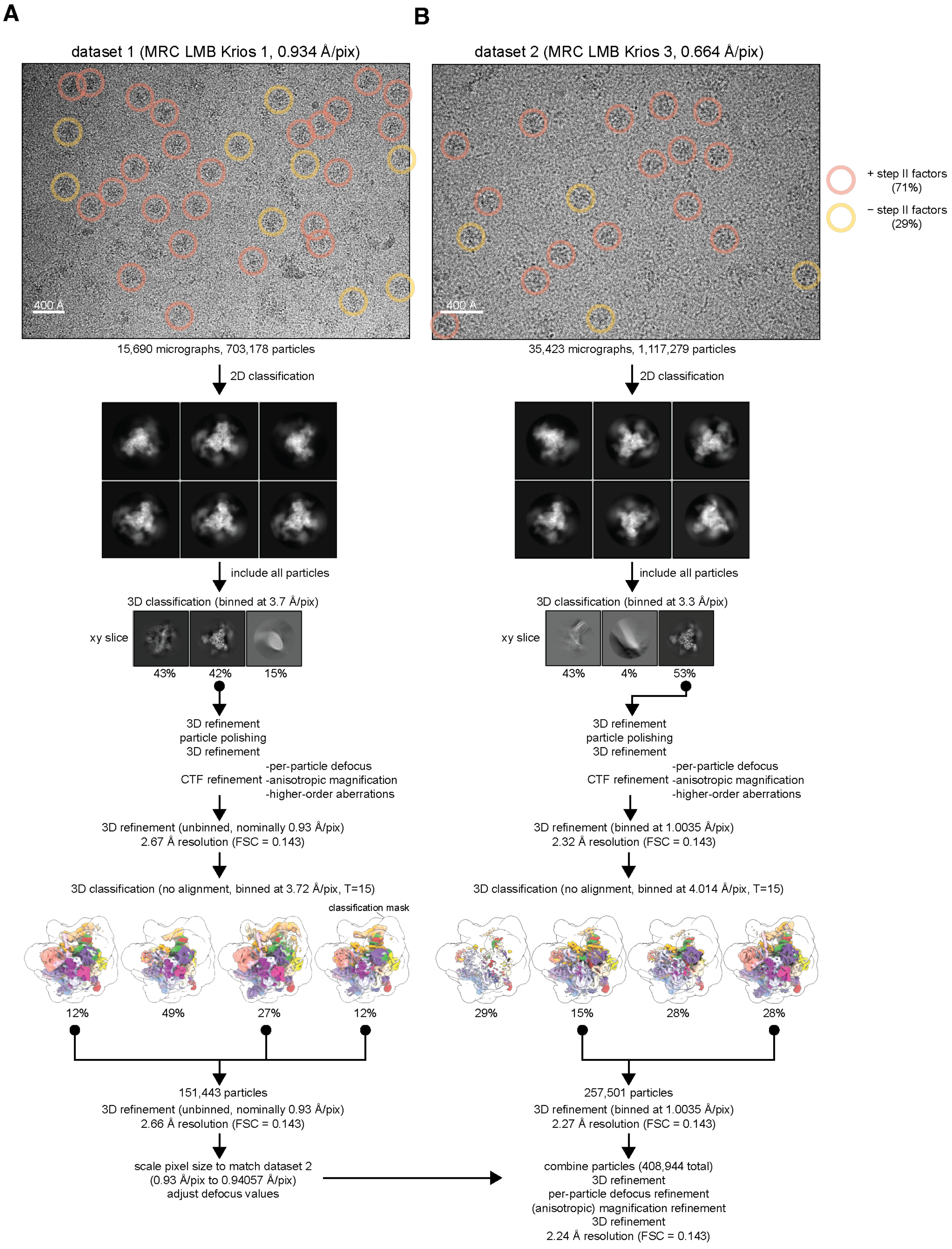
A
B
dataset 1 (MRC LMB Krios 1, 0.934 Å/pix)
dataset 2 (MRC LMB Krios 3, 0.664 Å/pix)
400 Å
400 Å
+ step II factors (71%)
− step II factors (29%)
15,690 micrographs, 703,178 particles
35,423 micrographs, 1,117,279 particles
2D classification
2D classification
include all particles
include all particles
3D classification (binned at 3.7 Å/pix)
3D classification (binned at 3.3 Å/pix)
xy slice
43%
42%
15%
xy slice
43%
4%
53%
3D refinement
particle polishing
3D refinement
CTF refinement -per-particle defocus
-anisotropic magnification
-higher-order aberrations
3D refinement
particle polishing
3D refinement
CTF refinement -per-particle defocus
-anisotropic magnification
-higher-order aberrations
3D refinement (unbinned, nominally 0.93 Å/pix)
2.67 Å resolution (FSC = 0.143)
3D refinement (binned at 1.0035 Å/pix)
2.32 Å resolution (FSC = 0.143)
3D classification (no alignment, binned at 3.72 Å/pix, T=15)
3D classification (no alignment, binned at 4.014 Å/pix, T=15)
classification mask
12%
49%
27%
12%
29%
15%
28%
28%
151,443 particles
3D refinement (unbinned, nominally 0.93 Å/pix)
2.66 Å resolution (FSC = 0.143)
257,501 particles
3D refinement (binned at 1.0035 Å/pix)
2.27 Å resolution (FSC = 0.143)
scale pixel size to match dataset 2
(0.93 Å/pix to 0.94057 Å/pix)
adjust defocus values
combine particles (408,944 total)
3D refinement
per-particle defocus refinement
(anisotropic) magnification refinement
3D refinement
2.24 Å resolution (FSC = 0.143)

**Figure 3-figure supplement 1. Spliceosome cryoEM data collection and general processing**. **A)** Example micrograph for dataset 1 with particles in states I or II circled in red and particles in state III circled in yellow. Example 2D classes are shown below, but 2D classification was not used for selection of particles for further processing. **B)** as **A)** but for dataset 2.

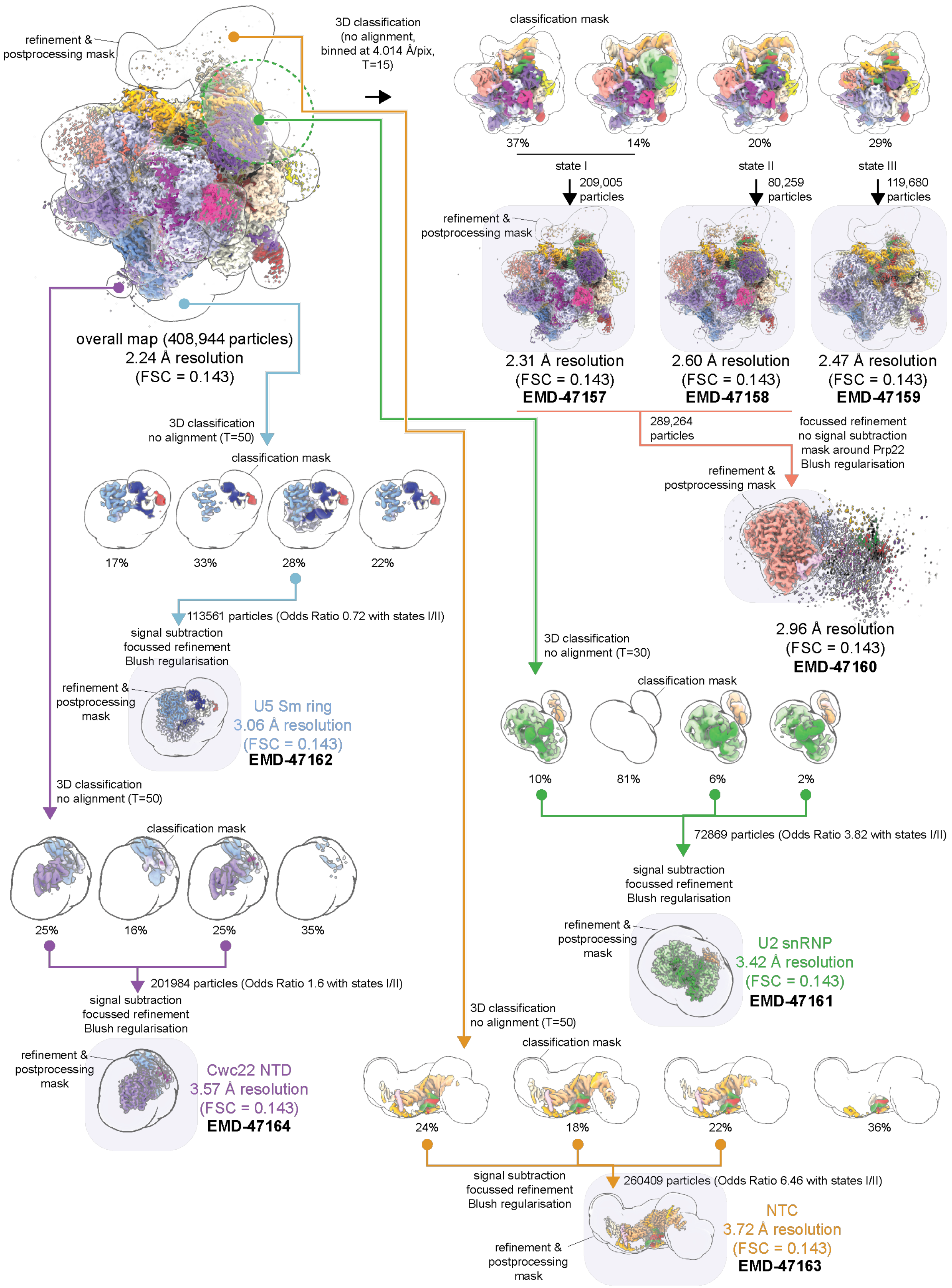

refinement & postprocessing mask
3D classification (no alignment, binned at 4.014 Å/pix, T=15)
classification mask
37%
14%
20%
29%
state I
state II
state III
209,005 particles
80,259 particles
119,680 particles
refinement & postprocessing mask
overall map (408,944 particles)
2.24 Å resolution
(FSC = 0.143)
2.31 Å resolution
(FSC = 0.143)
EMD-47157
2.60 Å resolution
(FSC = 0.143)
EMD-47158
2.47 Å resolution
(FSC = 0.143)
EMD-47159
289,264 particles
focussed refinement
no signal subtraction
mask around Prp22
Blush regularisation
3D classification
no alignment (T=50)
classification mask
refinement & postprocessing mask
17%
33%
28%
22%
2.96 Å resolution
(FSC = 0.143)
EMD-47160
113561 particles (Odds Ratio 0.72 with states I/II)
signal subtraction
focussed refinement
Blush regularisation
3D classification
no alignment (T=30)
refinement & postprocessing mask
U5 Sm ring
3.06 Å resolution
(FSC = 0.143)
EMD-47162
classification mask
10%
81%
6%
2%
3D classification
no alignment (T=50)
72869 particles (Odds Ratio 3.82 with states I/II)
classification mask
signal subtraction
focussed refinement
Blush regularisation
25%
16%
25%
35%
refinement & postprocessing mask
U2 snRNP
3.42 Å resolution
(FSC = 0.143)
EMD-47161
201984 particles (Odds Ratio 1.6 with states I/II)
signal subtraction
focussed refinement
Blush regularisation
3D classification
no alignment (T=50)
refinement & postprocessing mask
Cwc22 NTD
3.57 Å resolution
(FSC = 0.143)
EMD-47164
classification mask
24%
18%
22%
36%
signal subtraction
focussed refinement
Blush regularisation
260409 particles (Odds Ratio 6.46 with states I/II)
refinement & postprocessing mask
NTC
3.72 Å resolution
(FSC = 0.143)
EMD-47163

**Figure 3-figure supplement 2. Focused classification and refinement scheme for regions of P complex**. Final maps deposited to the EMDB are highlighted.

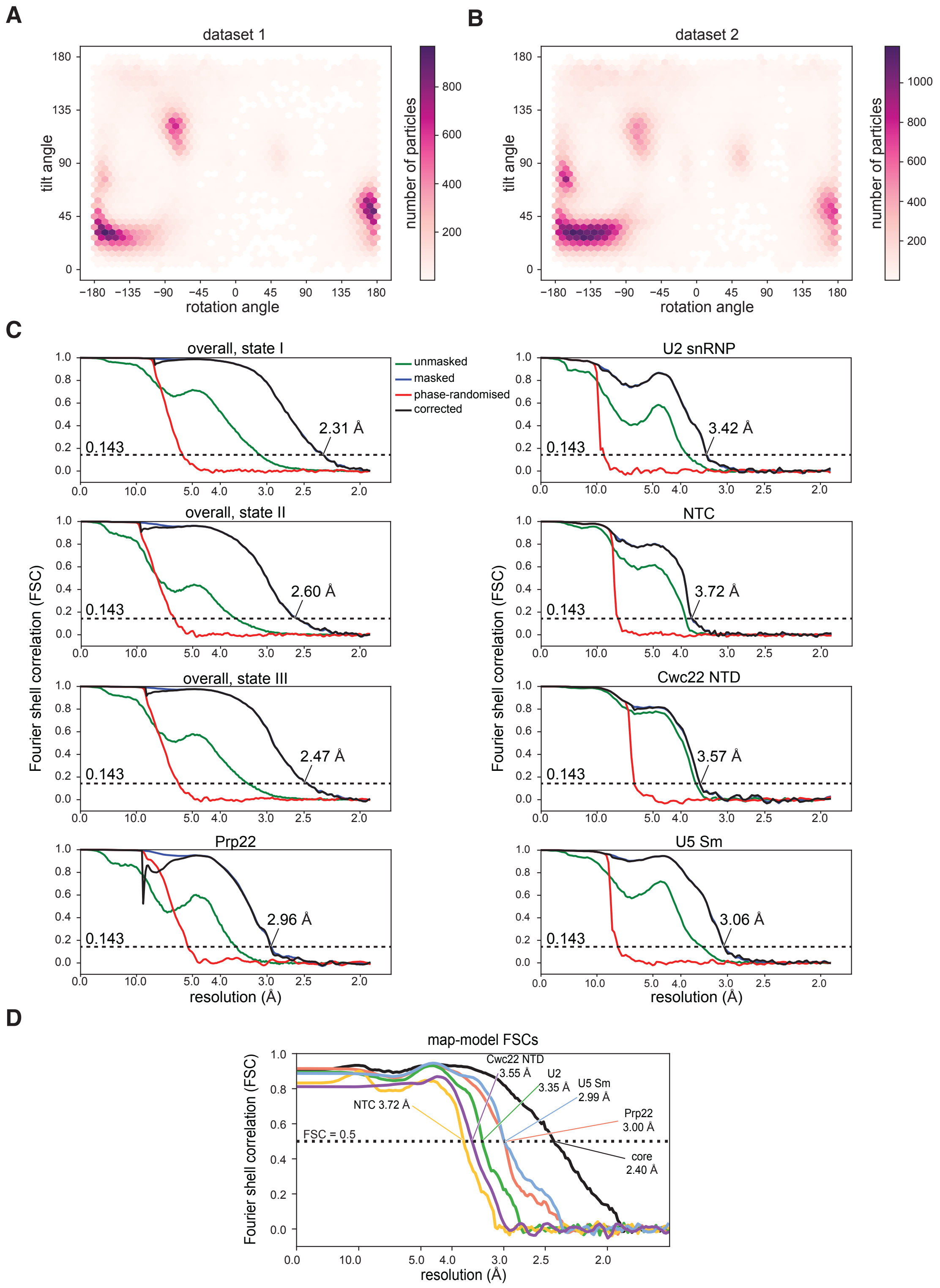


**Figure 3-figure supplement 3. A,B)** Orientation distribution plot for state I separated by datasets. **C)** Gold-standard Fourier-Shell Correlation curves for the overall reconstructions and focus-refined maps. **D)** Map-model Fourier-Shell Correlation, calculated using PHENIX.

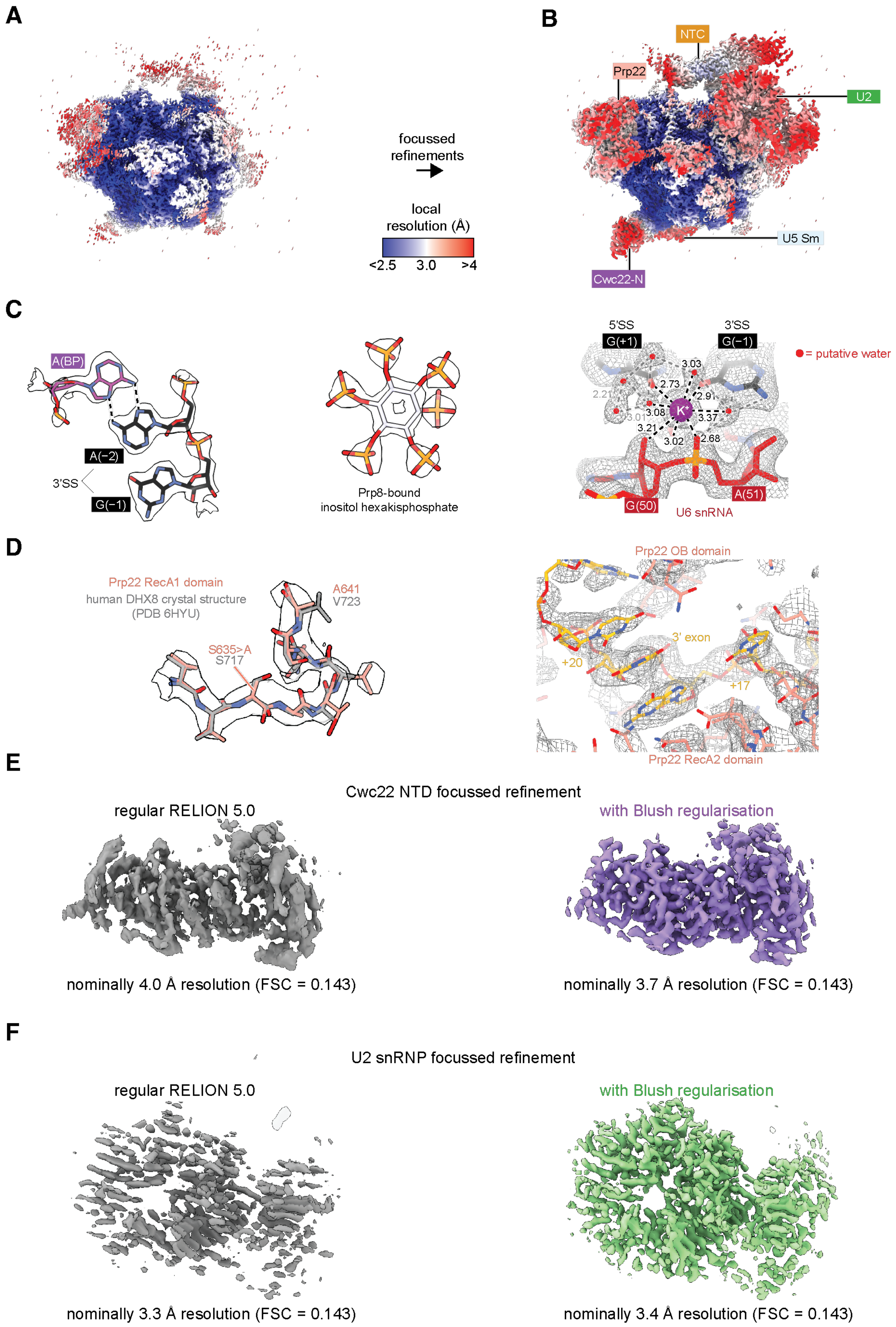

A
B
focussed
refinements
local
resolution (Å)
<2.5
3.0
>4
NTC
Prp22
U2
U5 Sm
Cwc22-N
C
A(BP)
A(−2)
3'SS
G(−1)
Prp8-bound
inositol hexakisphosphate
5'SS
G(+1)
3'SS
G(−1)
= putative water
3.03
2.73
2.21
2.91
3.01
3.08
3.37
3.21
3.02
2.68
K+
G(50)
A(51)
U6 snRNA
D
Prp22 RecA1 domain
human DHX8 crystal structure
(PDB 6HYU)
A641
V723
S635>A
S717
Prp22 OB domain
3' exon
+20
+17
Prp22 RecA2 domain
E
Cwc22 NTD focussed refinement
regular RELION 5.0
with Blush regularisation
nominally 4.0 Å resolution (FSC = 0.143)
nominally 3.7 Å resolution (FSC = 0.143)
F
U2 snRNP focussed refinement
regular RELION 5.0
with Blush regularisation
nominally 3.3 Å resolution (FSC = 0.143)
nominally 3.4 Å resolution (FSC = 0.143)

**Figure 3-figure supplement 4. A)** The overall P complex reconstruction (state I) colored by local resolution as calculated within RELION. **B)** Local resolution after focused refinements, colored as in **A**). **C)** Representative density for the P complex core. **D)** (Left) Density for the loop containing the S635A dominant-negative mutation in Prp22. The model is superimposed over the crystal structure of DHX8 (human Prp22) in the same region (PDB ID 6HYU; Felisberto-Rodrigues et al., 2019). (Rigth) Density for the 3' exonic RNA within Prp22. **E)** Focused refinement of the Cwc22 NTD without and with Blush regularization. All refinement settings and inputs were the same except for the usage of Blush. Only dataset 2 particles were used for this comparison. **F)** As for **E)** but with the U2 snRNP. Note that the resolution estimate is inflated without Blush regularization.

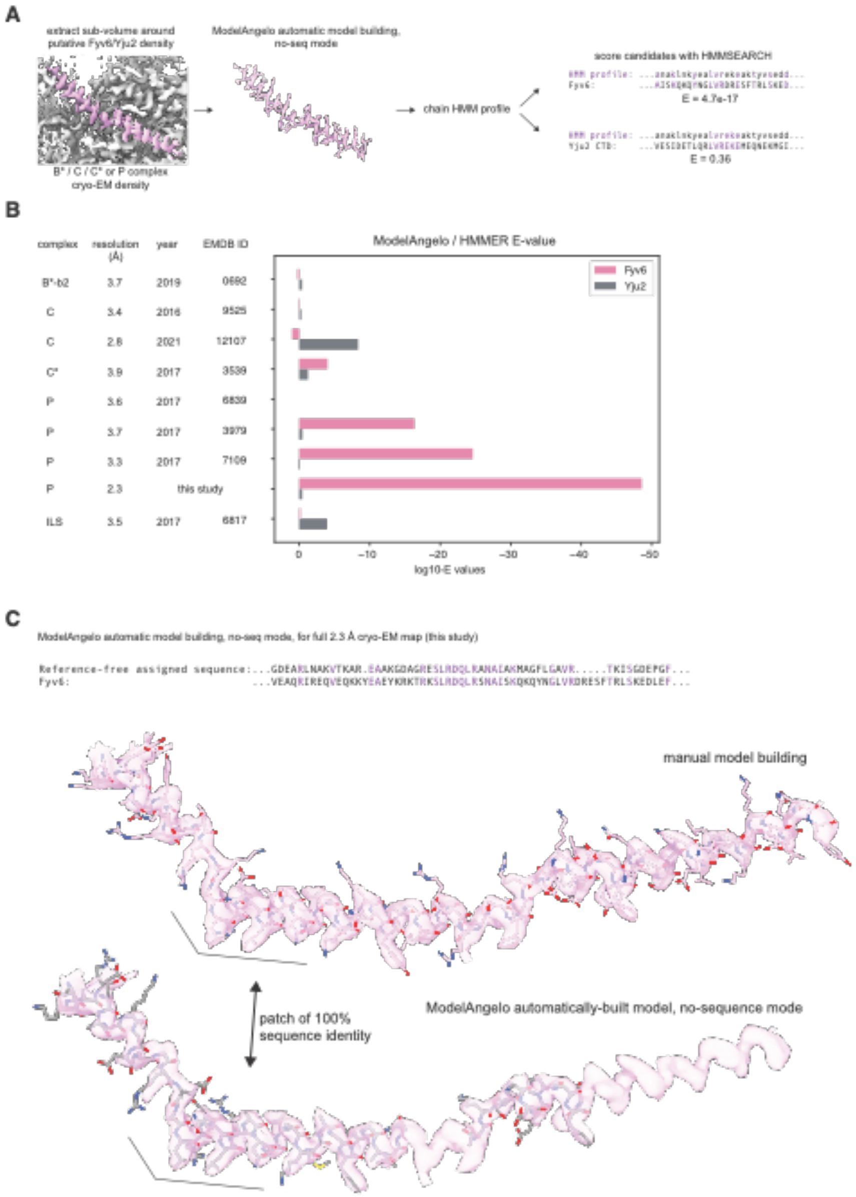


**Figure 4-figure supplement 1. Comparison of fits to density of Yju2 versus Fyv6 in each complex. A)** Cryo-EM densities were boxed around Fyv6 or Yju2 using ChimeraX. These were used as inputs to ModelAngelo operated in “no-seq” mode. The HMM profiles for the chains built into the putative Fyv6/Yju2 density were used to score all spliceosomal proteins using hmmsearch. **B)** Expectation values (E-values) for hmmsearch against Fyv6 or Yju2 protein sequences. Lower E-values indicate a better match to the profile. **C)** Comparison of the model built by ModelAngelo in no-seq mode to the manually built model for Fyv6 in P complex State I (this study). Despite not providing the sequence for Fyv6, several patches had 100% sequence identity.

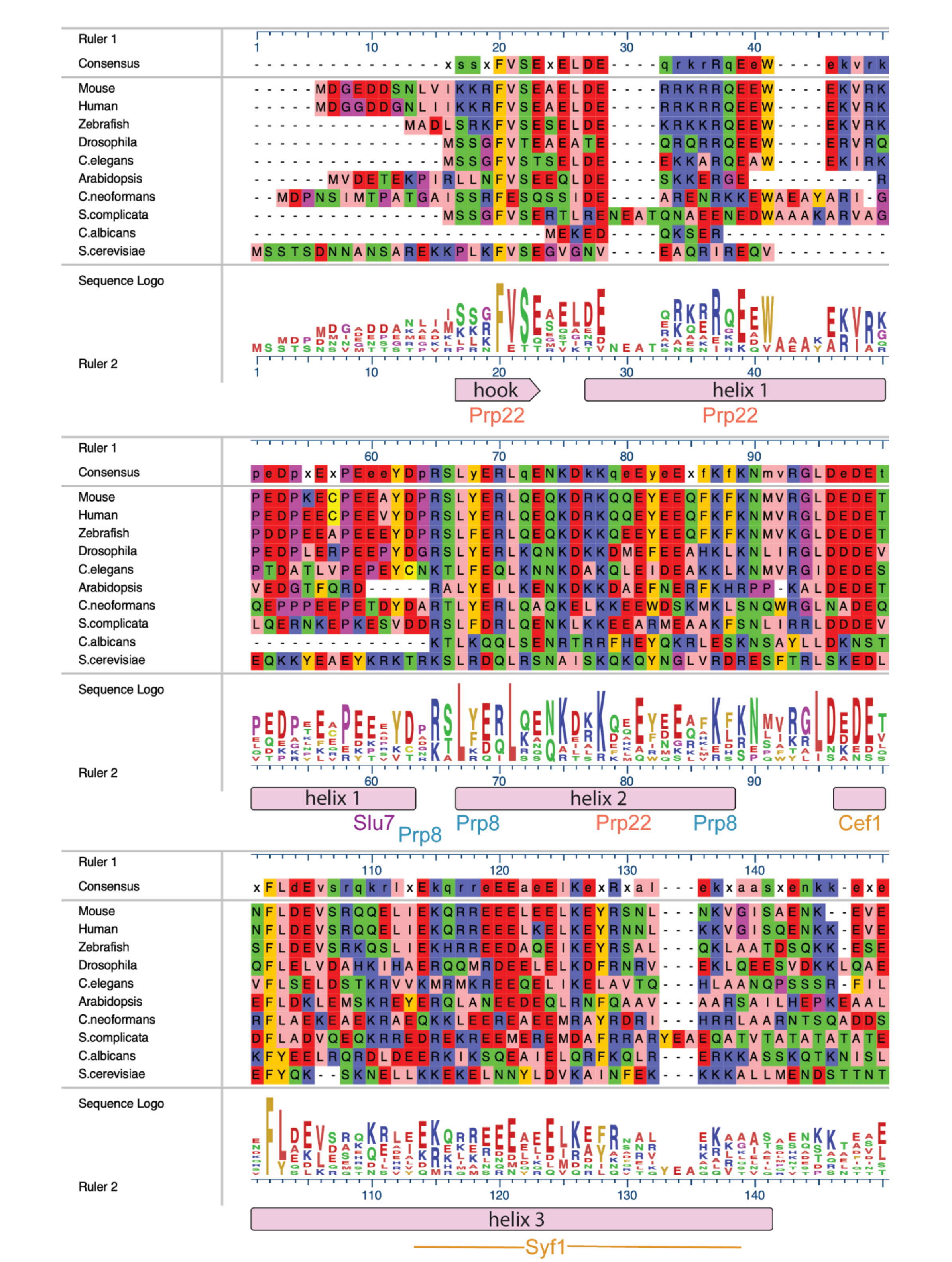


**Figure 4-figure supplement 2 (continued next page)**

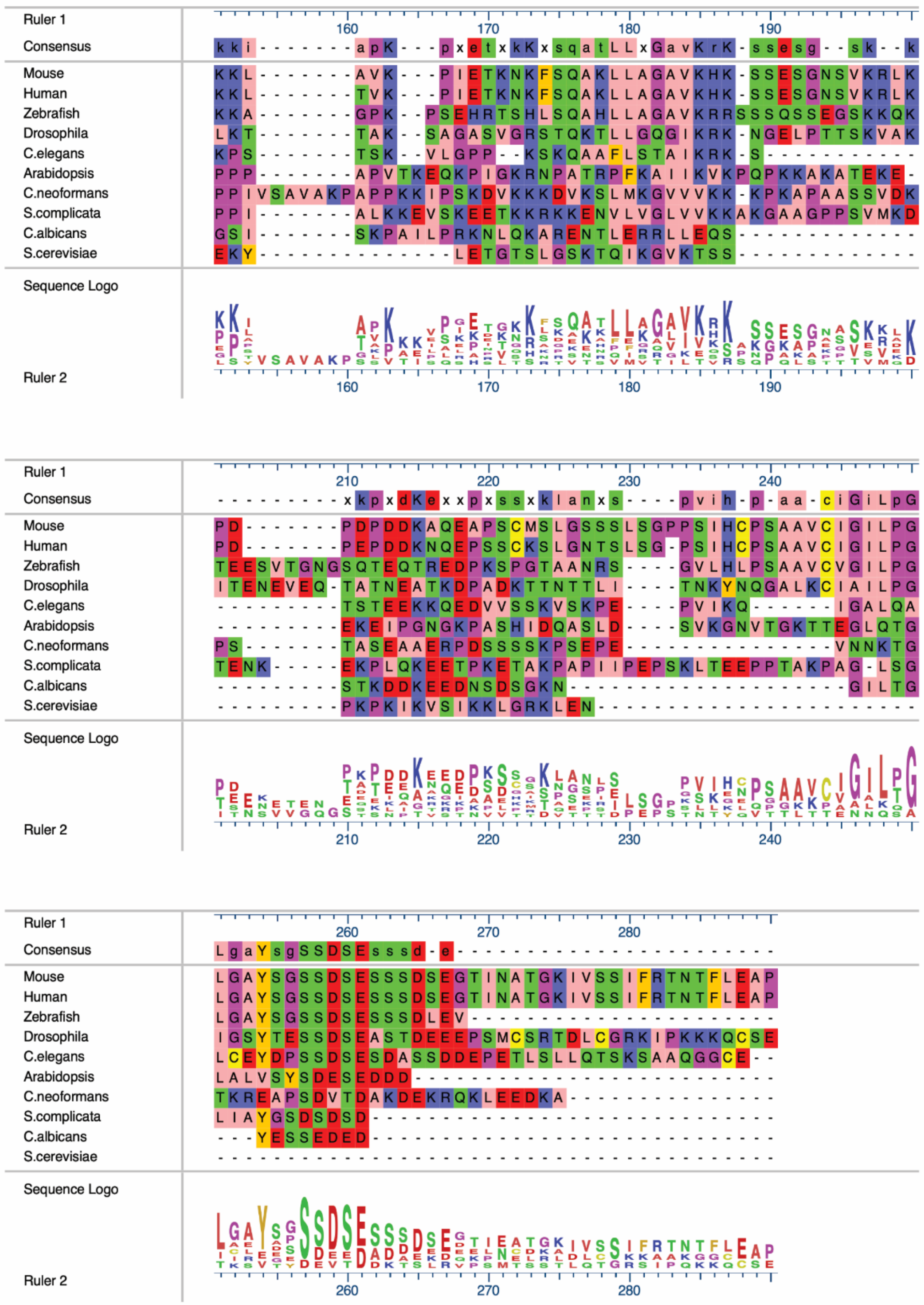

**Figure 4-figure supplement 2. Alignments of Fyv6 homolog protein sequences.** Sequence-based alignment of Fyv6 and homologs from other model organisms. Alignment, consensus, and sequence logos are shown. Regions of Fyv6 protein structure determined from cryo-EM model are annotated below corresponding sequence. Sequences were aligned and visualized using MegAlign Pro (DNAStar) and Clustal Omega.

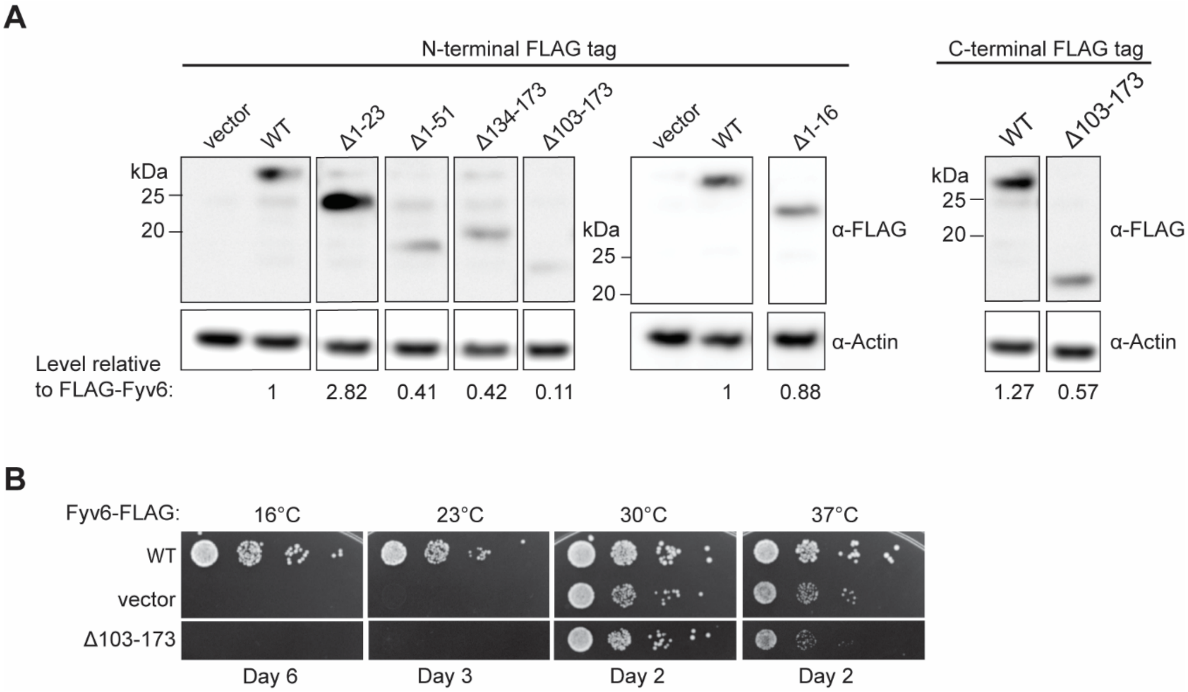


**Figure 5-figure supplement 1. A)** Expression of FLAG-tagged Fyv6 truncations visualized by western blot using anti-FLAG antibody. Actin was used as a loading control. **B)** Temperature growth assay of C-terminally FLAG-tagged WT Fyv6 and the Δ103-173 truncation on –Trp plates. Plates were imaged after days indicated.

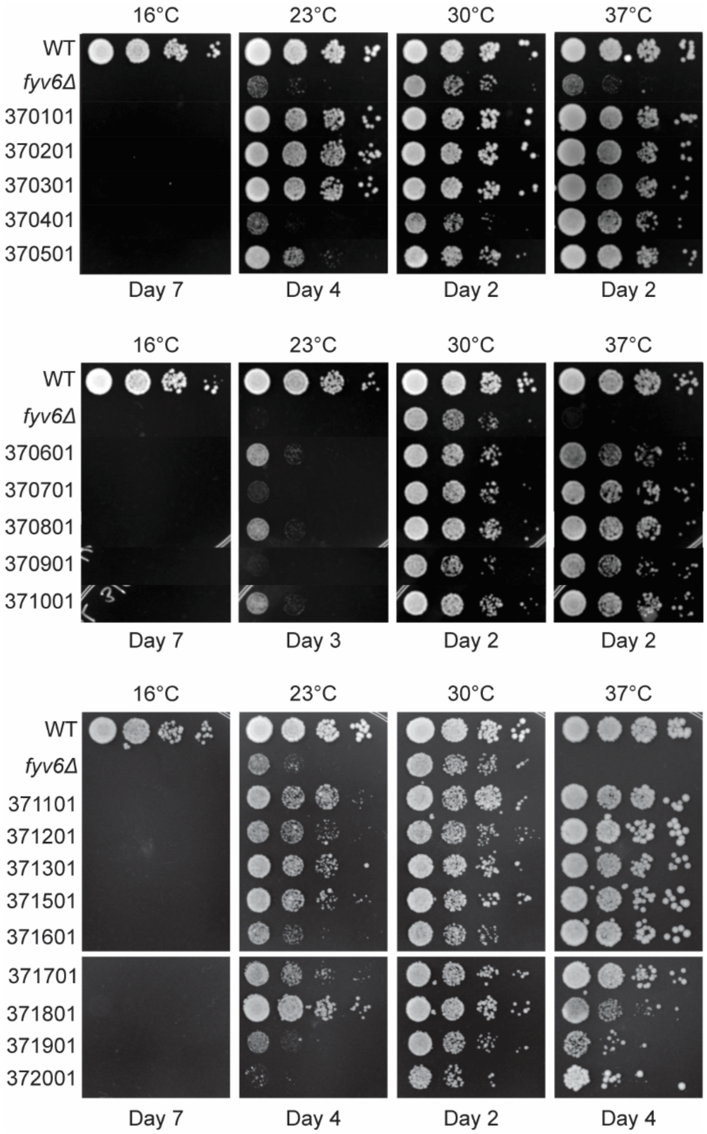


**Figure 6-figure supplement 1. Isolated suppressor strains for *fyv6Δ* grown at different temperatures.** Strains were grown on YPD plates and imaged after the number of days indicated.

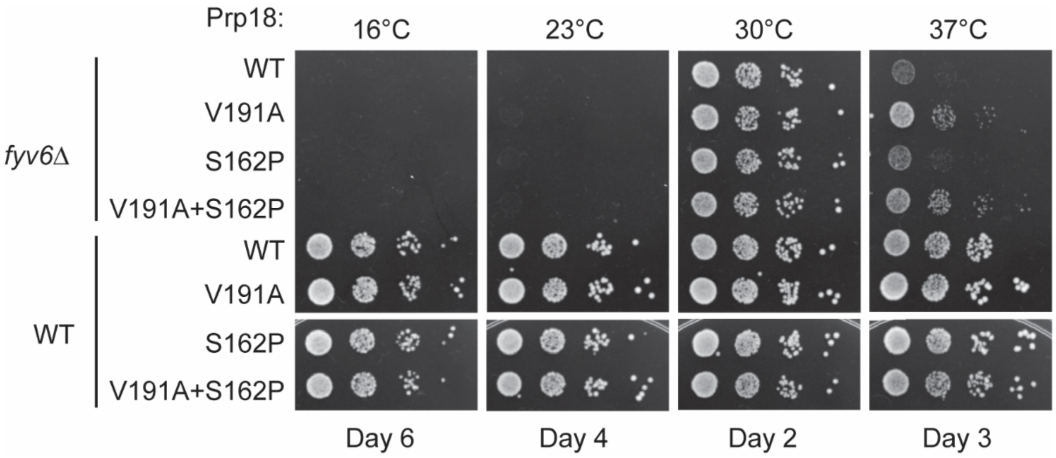


**Figure 6-figure supplement 2. Genetic interactions between Prp18 and Fyv6.** Spot dilution temperature growth assay for genetic interactions of *FYV6* deletion with Prp18 alleles on –Ura DO plates. Plate images were taken on days indicated.

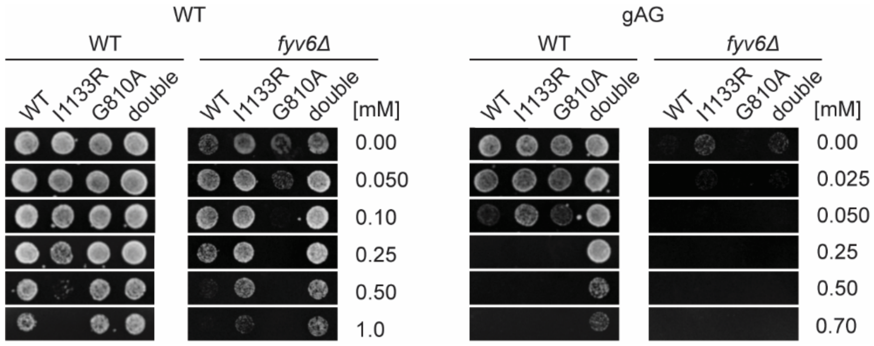


**Figure 7-figure supplement 1. ACT1-CUP1 plate images for the data shown in Figure 7H.** Images of yeast growth on copper-containing –Leu DO media shown after 48 (WT) or 72 h (*fyv6Δ*) for strains containing the indicated ACT1-CUP1 reporters.

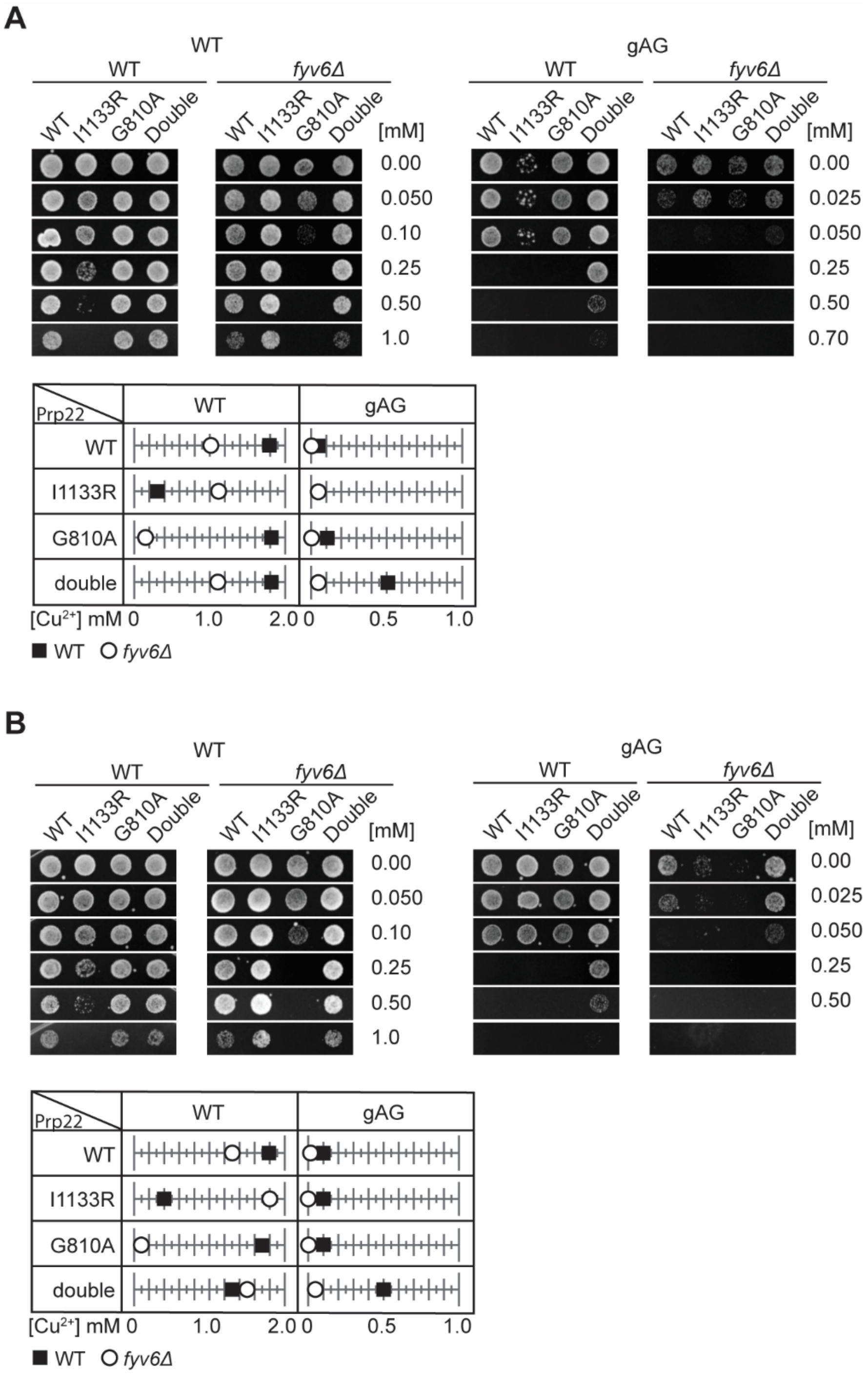


**Figure 7-figure supplement 2. Replicate ACT1-CUP1 assays to those shown in Figure 7. A,B)** Plate images and quantitation of copper tolerances for two additional replicates of the assay shown in Fig. 7G. Images of representative yeast growth on copper-containing –Leu D.O. media shown after 48 (WT) or 72 h (*fyv6Δ*) for strains containing the indicated ACT1-CUP1 reporters.

### Supplementary Table 1: STAR Mapping Statistics for samples with UPF1

See associated Excel file for this table.

**Supplementary Table 2. RNA-seq datasets used for analysis.**

| Datasets | Description/Encode # | Figures |
|---|---|---|
| WT-16-12-8, WT-16-1-5, WT-30-12-8, WT-30-1-5, Fyv6-16-12-8, Fyv6-16-1-5, Fyv6-30-12-8, Fyv6-30-1-5, Fyv6-37-12-8, Fyv6-37-1-5 | Temperature shifted RNA-seq datasets for WT *upf1Δ* and *fyv6Δ upf1Δ* yeast strains | Fig. 1 |
| WT-0409, WT-0417, Fyv6-0409, Fyv6-0417 | RNA-seq datasets for WT and fyv6Δ yeast strains | Fig. S1C |
| WT0803, WT0811, Del0803, Del0811 | RNA-seq datasets for WT upf1Δ and fyv6Δ upf1Δ yeast strains | Fig. S1A,D; Fig. 2 |

**Supplementary Table 3. Sequences of introns between the branch point and 3' SS in ACT1-CUP1 reporters.**

| BP-3' SS distance | Intron sequence between branch site and 3' SS* |
|---|---|
| 9 nt | **UACUAACA**UCGAUUAUA**UAG** |
| 12 nt | **UACUAACA**UCGAUUUGUUUA**UAG** |
| 15 nt | **UACUAACA**UCGAUUAUAUGUUUA**UAG** |
| 21 nt | **UACUAACA**UCGUUCUUCUUUCCGAUUAUA**UAG** |
| 27 nt | **UACUAACA**UCGUUCUUCUUUCCGAUUAUAUGUUUA**UAG** |
| 38 nt | **UACUAACA**UCGAUUGCUUCAUUCUUUUUGUUGCUAUAUUAUAUGUU**UAG** |
| 42 nt | **UACUAACA**UCGAAACAUUGCUUCAUUCUUUUUGUUGCUAUAUUAUAUGUU **UAG** |
| 46 nt | **UACUAACA**UCGAAACAACAAUUGCUUCAUUCUUUUUGUUGCUAUAUUAUA UGUU**UAG** |
| 50 nt | **UACUAACA**UCGAAACAACAAACGAUUGCUUCAUUCUUUUUGUUGCUAUAU UAUAUGUU**UAG** |

*Branch site and 3' SS sequences in bold

### Supplementary Table 4. Cryo-EM data processing, refinement, and validation statistics

See associated Excel file for this table.

**Supplementary Table 5. Mutations identified in *fyv6Δ* suppressor strains selected at 37°C**

| Gene | Chromosome | Position | Substitution | Mutation | Strain(s) | Human Gene | Human Residue |
|---|---|---|---|---|---|---|---|
| PRP8 | VIII | 432198 | G->T | S1584Y | 370201 | | S1512 |
| | | 432198 | G->A | S1584F*^ | 372003 | | S1512 |
| | | 431365 | C->G | V1862L | 370801 | | I1790 |
| | | 431347 | C->T | G1868R | 370701 | | G1796 |
| | | 431004 | G->C | T1982S^ | 370501 | | T1910 |
| SLU7 | IV | 619621 | C->T | E9K | 370301 | SLU7 | N/A |
| | | 619576 | C->T | A24R** | 370101 | | N/A |
| | | 619575 | G->C | | | | |
| | | 619570 | C->T | E26K** | | | N/A |
| | | 619562 | A->C | N28K** | | | D58 |
| CDC40 (PRP17) | IV | 1203588 | G->A | P208L | 371102 | CDC40 | A245 |
| | | 1203459 | G->A | A251V | 370601 | | Y367 |
| | | 1203266 | C->A | K315N | 371701 | | K450 |
| | | 1203055 | C->A | G386W | 370901 | | A511 |
| Cef1 | XIII | 693490 | C->T | A37V | 370401 | CDC5L | A35 |
| | | 693905 | G->T | M175I | 371201, 371302 | | Q175 |
| | | 693958 | A->C | Q193P | 370804 | | A193 |
| CLF1 (SYF3) | XII | 384437 | G->A | T33I^^ | 371001 | CRNKL1 | P210 |
| RSE1 | XIII | 175909 | A->T | D799E*, ***,**** | 371601, 371901, 372003 | SF3B3 | N/A |
| | | 175912 | G->T | D798E*** | 371901 | | N/A |
| | | 175899 | T->C | K803E*** | | | N/A |
| | | 175896 | C->T | E804K*** | | | N/A |
| | | 175883 | A->T | I808K*** | | | N/A |
| LSR1 | II | 681838 | T->C | A25G | 371802 | RNU2 | A24 |
| PRP22 | V | 182238 | T->G | I1133R** | 371601 | DHX8 | N/A |

*Mutations arose together in strain 372003. **Mutations arose together in strain 370101. ***Mutations arose together in strain 371901. ****Mutations arose together in 371601. ^Second-strongest suppressors of *FYV6* deletion based on colony size. ^^Strongest suppressor of *FYV6* deletion based on colony size.

**Supplementary Table 6. Yeast strains used in this study.**

| Strain name | Genotype | Description | Source |
|---|---|---|---|
| yAAH0434 | MATα cup1Δ ura3 his3 trp1 lys2 ade2 leu2 | $Cu^{2+}$ sensitive strain | David Brow |
| yAAH3353 | yAAH0434 + fyv6Δ::hphMX | $Cu^{2+}$ sensitive *fyv6Δ* strain | Lipinski et al., 2023 |
| yAAH3399 | yAAH0434+upf1Δ::KanMX | $Cu^{2+}$ sensitive *upf1Δ* strain | Lipinski et al., 2023 |
| yAAH3400 | yAAH3353+upf1Δ::KanMX | $Cu^{2+}$ sensitive *fyv6Δ upf1Δ* strain | Lipinski et al., 2023 |
| yAAH3403 | yAAH0434 + dbr1Δ::NatR | $Cu^{2+}$ sensitive *dbr1Δ* strain | This study |
| yAAH3404 | yAAH3353 + dbr1Δ::NatR | $Cu^{2+}$ sensitive *fyv6Δ dbr1Δ* strain | This study |
| yAAH3388 | yAAH3353 + pAAH1555 | $Cu^{2+}$ sensitive Fyv6 shuffle strain | This study |
| yAAH3393 | yAAH3353 + pAAH0135 | vector control, contains pRS414 | This study |
| yAAH3419 | yAAH3353 + pAAH1572 | FLAG-Fyv6 (N-terminal tag) | This study |
| yAAH3434 | yAAH3353 + pAAH1577 | FLAG-Fyv6-Δ1-16 | This study |
| yAAH3435 | yAAH3353 + pAAH1578 | FLAG-Fyv6-Δ1-23 | This study |
| yAAH3436 | yAAH3353 + pAAH1579 | FLAG-Fyv6-Δ1-51 | This study |
| yAAH3437 | yAAH3353 + pAAH1580 | FLAG-Fyv6-Δ134-173 | This study |
| yAAH3438 | yAAH3353 + pAAH1581 | FLAG-Fyv6-Δ103-173 | This study |
| yAAH3418 | yAAH3353 + pAAH1573 | Fyv6-FLAG (C-terminal tag) | This study |
| yAAH3442 | yAAH3353 + pAAH1586 | Fyv6-Δ103-173 -FLAG | This study |
| yAAH3455 | yAAH0434 + pAAH1602 | $Prp18^{WT}$ merodiploid | This study |
| yAAH3456 | yAAH0434 + pAAH1603 | $Prp18^{WT}/Prp18^{V191A}$ merodiploid | This study |
| yAAH3457 | yAAH0434 + pAAH1604 | $Prp18^{WT}$ /$Prp18^{S162P}$ merodiploid | This study |
| yAAH3458 | yAAH0434 + pAAH1605 | $Prp18^{WT}$ /$Prp18^{S162P+V191A}$ merodiploid | This study |
| yAAH3459 | yAAH3353 + pAAH1602 | $Prp18^{WT}$ merodiploid in *fyv6Δ* strain | This study |
| yAAH3460 | yAAH3353 + pAAH1603 | $Prp18^{WT}/Prp18^{V191A}$ merodiploid in *fyv6Δ* strain | This study |
| yAAH3461 | yAAH3353 + pAAH1604 | $Prp18^{WT}$ /$Prp18^{S162P}$ merodiploid in *fyv6Δ* strain | This study |
| yAAH3462 | yAAH3353 + pAAH1605 | $Prp18^{WT}$ /$Prp18^{S162P+V191A}$ merodiploid in *fyv6Δ* strain | This study |
| yAAH3405 | yAAH3403 + pAAH0470 | *dbr1Δ* + WT ACT1-CUP1 (38 nt) | This study |
| yAAH3410 | yAAH3404 + pAAH0470 | *dbr1Δ fyv6Δ* + WT ACT1-CUP1 (38 nt) | This study |
| yAAH3498 | yAAH3403 + pAAH1632 | *dbr1Δ* + ACT1-CUP1 (9 nt) | This study |
| yAAH3499 | yAAH3403 + pAAH1633 | *dbr1Δ* + ACT1-CUP1 (12 nt) | This study |
| yAAH3500 | yAAH3403 + pAAH1634 | *dbr1Δ* + ACT1-CUP1 (21 nt) | This study |
| yAAH3501 | yAAH3403 + pAAH1635 | *dbr1Δ* + ACT1-CUP1 (27 nt) | This study |
| yAAH3503 | yAAH3404 + pAAH1633 | *dbr1Δ fyv6Δ* + ACT1-CUP1 (9 nt) | This study |

| | | | |
|---|---|---|---|
| yAAH3504 | yAAH3404 + pAAH1634 | *dbr1Δ fyv6Δ* + ACT1-CUP1 (12 nt) | This study |
| yAAH3505 | yAAH3404 + pAAH1635 | *dbr1Δ fyv6Δ* + ACT1-CUP1 (21 nt) | This study |
| yAAH3506 | yAAH3404 + pAAH1632 | *dbr1Δ fyv6Δ* + ACT1-CUP1 (27 nt) | This study |
| yAAH3509 | yAAH3403 + pAAH1636 | *dbr1Δ* + ACT1-CUP1 (15 nt) | This study |
| yAAH3510 | yAAH3403 + pAAH1637 | *dbr1Δ* + ACT1-CUP1 (42 nt) | This study |
| yAAH3511 | yAAH3403 + pAAH1638 | *dbr1Δ* + ACT1-CUP1 (46 nt) | This study |
| yAAH3512 | yAAH3403 + pAAH1639 | *dbr1Δ* + ACT1-CUP1 (50 nt) | This study |
| yAAH3513 | yAAH3404 + pAAH1636 | *dbr1Δ fyv6Δ* + ACT1-CUP1 (15 nt) | This study |
| yAAH3514 | yAAH3404 + pAAH1637 | *dbr1Δ fyv6Δ*+ ACT1-CUP1 (42 nt) | This study |
| yAAH3515 | yAAH3404 + pAAH1638 | *dbr1Δ fyv6Δ*+ ACT1-CUP1 (46 nt) | This study |
| yAAH3516 | yAAH3404 + pAAH1639 | *dbr1Δ fyv6Δ*+ ACT1-CUP1 (50 nt) | This study |
| yAAH3517 | yAAH0434+syf1Δ::KanMX + pAAH1624 | $Cu^{2+}$ sensitive Syf1 shuffle strain | This study |
| yAAH3518 | yAAH3517+ fyv6Δ::HygR | $Cu^{2+}$ sensitive Syf1 shuffle strain with *fyv6Δ* | This study |
| yAAH3519 | yAAH0434+syf1Δ::KanMX + pAAH1625 | WT Syf1 | This study |
| yAAH3577 | yAAH0434+syf1Δ::KanMX + +pAAH1666 | Syf1 Δ817-859 | This study |
| yAAH3578 | yAAH0434+syf1Δ::KanMX + fyv6Δ::HygR + pAAH1666 | *fyv6Δ* + Syf1 Δ817-859 | This study |
| yAAH3579 | yAAH0434+syf1Δ::KanMX + +pAAH1667 | Syf1 Δ778-859 | This study |
| yAAH3580 | yAAH0434+syf1Δ::KanMX + fyv6Δ::HygR + pAAH1667 | *fyv6Δ* + Syf1 Δ778-859 | This study |
| yAAH3581 | yAAH0434+syf1Δ::KanMX + +pAAH1668 | Syf1 Δ634-859 | This study |
| yAAH3582 | yAAH0434+syf1Δ::KanMX + fyv6Δ::HygR + pAAH1668 | *fyv6Δ* + Syf1 Δ634-859 | This study |
| yAAH3583 | yAAH0434+syf1Δ::KanMX + fyv6Δ::HygR + pAAH1625 (WT Syf1) | *fyv6Δ* + WT Syf1 | This study |
| yAAH3593 | yAAH0434 + cef1Δ::KanMX + pAAH1658 | $Cu^{2+}$ sensitive Cef1 shuffle strain | This study |
| yAAH3594 | yAAH3353 + cef1Δ::KanMX + pAAH1658 | $Cu^{2+}$ sensitive Cef1 shuffle strain with fyv6Δ | This study |
| yAAH3634 | yAAH0434 + cef1Δ::KanMX + pAAH1611 | $Cef1^{WT}$ | This study |
| yAAH3636 | yAAH0434 + cef1Δ::KanMX + pAAH1642 | $Cef1^{M175I}$ | This study |
| yAAH3637 | yAAH0434 + cef1Δ::KanMX + pAAH1612 | $Cef1^{A37P}$ | This study |
| yAAH3638 | yAAH0434 + cef1Δ::KanMX + pAAH1641 | $Cef1^{A37V}$ | This study |
| yAAH3639 | yAAH0434 + cef1Δ::KanMX + pAAH1614 | $Cef1^{V36R}$ | This study |

| | | | |
|---|---|---|---|
| yAAH3640 | yAAH0434 + cef1Δ::KanMX + pAAH1613 | $Cef1^{S48R}$ | This study |
| yAAH3641 | yAAH0434 + cef1Δ::KanMX + pAAH1643 | $Cef1^{Q193P}$ | This study |
| yAAH3655 | yAAH3353 + cef1Δ::KanMX + pAAH1612 | *fyv6Δ* $Cef1^{A37P}$ | This study |
| yAAH3656 | yAAH3353 + cef1Δ::KanMX + pAAH1642 | *fyv6Δ* $Cef1^{M175I}$ | This study |
| yAAH3657 | yAAH3353 + cef1Δ::KanMX + pAAH1641 | *fyv6Δ* $Cef1^{A37V}$ | This study |
| yAAH3658 | yAAH3353 + cef1Δ::KanMX + pAAH1614 | *fyv6Δ* $Cef1^{V36R}$ | This study |
| yAAH3659 | yAAH3353 + cef1Δ::KanMX + pAAH1613 | *fyv6Δ* $Cef1^{S48R}$ | This study |
| yAAH3660 | yAAH3353 + cef1Δ::KanMX + pAAH1643 | *fyv6Δ* $Cef1^{Q193P}$ | This study |
| yAAH3661 | yAAH3353 + cef1Δ::KanMX + pAAH1658 | *fyv6Δ* $Cef1^{WT}$ | This study |
| yAAH0117 | ade2, cup1Δ:ura3 his3 leu2 lys2 prp8Δ:lys2 trp1 pJU169:PRP8(URA) | $Cu^{2+}$ sensitive Prp8 shuffle strain | Christine Guthrie |
| yAAH3352 | yAAH117 + fyv6Δ::HygR | $Cu^{2+}$ sensitive *fyv6Δ* Prp8 shuffle strain | Lipinski et al., 2023 |
| yAAH3093 | ade2, cup1Δ:ura3 his3 leu2 lys2 prp8Δ:lys2 trp1 +pAAH1440 | $Prp8^{WT}$ | This study |
| yAAH3683 | ade2, cup1Δ:ura3 his3 leu2 lys2 prp8Δ:lys2 trp1 +pAAH1659 | $Prp8^{S1584Y}$ | This study |
| yAAH3684 | ade2, cup1Δ:ura3 his3 leu2 lys2 prp8Δ:lys2 trp1 +pAAH1660 | $Prp8^{S1584F}$ | This study |
| yAAH3685 | ade2, cup1Δ:ura3 his3 leu2 lys2 prp8Δ:lys2 trp1 +pAAH1661 | $Prp8^{V1862L}$ | This study |
| yAAH3686 | ade2, cup1Δ:ura3 his3 leu2 lys2 prp8Δ:lys2 trp1 +pAAH1662 | $Prp8^{G1868R}$ | This study |
| yAAH3687 | ade2, cup1Δ:ura3 his3 leu2 lys2 prp8Δ:lys2 trp1 +pAAH1663 | $Prp8^{T1982S}$ | This study |
| yAAH3688 | ade2, cup1Δ:ura3 his3 leu2 lys2 prp8Δ:lys2 trp1 fyv6Δ::HygR +pAAH1659 | *fyv6Δ* $Prp8^{S1584Y}$ | This study |
| yAAH3689 | ade2, cup1Δ:ura3 his3 leu2 lys2 prp8Δ:lys2 trp1 fyv6Δ::HygR +pAAH1660 | *fyv6Δ* $Prp8^{S1584F}$ | This study |
| yAAH3690 | ade2, cup1Δ:ura3 his3 leu2 lys2 prp8Δ:lys2 trp1 fyv6Δ::HygR +pAAH1661 | *fyv6Δ* $Prp8^{V1862L}$ | This study |
| yAAH3691 | ade2, cup1Δ:ura3 his3 leu2 lys2 prp8Δ:lys2 trp1 fyv6Δ::HygR +pAAH1662 | *fyv6Δ* $Prp8^{G1868R}$ | This study |
| yAAH3692 | ade2, cup1Δ:ura3 his3 leu2 lys2 prp8Δ:lys2 trp1 fyv6Δ::HygR +pAAH1663 | *fyv6Δ* $Prp8^{T1982S}$ | This study |

| | | | |
|---|---|---|---|
| yAAH3693 | ade2, cup1Δ:ura3 his3 leu2 lys2 prp8Δ:lys2 trp1 fyv6Δ::HygR +pAAH1440 | *fyv6Δ* $Prp8^{WT}$ | This study |
| yAAH1930 | MATa ade2 cup1Δ::ura3 his3 leu2 lys2 trp1 ura3 GAL+ prp22Δ::loxP pPrp22 (URA3) | Prp22 Shuffle Strain | Charles Query |
| yAAH1931 | MATa ade2 cup1Δ::ura3 his3 leu2 lys2 trp1 ura3 GAL+ prp22Δ::loxP +pAAH1042 | $Prp22^{WT}$ | Charles Query |
| yAAH3377 | MATa ade2 cup1Δ::ura3 his3 leu2 lys2 trp1 ura3 GAL+ prp22Δ::loxP fyv6Δ::hygR pPrp22 (URA3) | *fyv6Δ* Prp22 Shuffle Strain | Lipinski et al., 2023 |
| yAAH3379 | MATa ade2 cup1Δ::ura3 his3 leu2 lys2 trp1 ura3 GAL+ prp22Δ::loxP fyv6Δ::hygR + pAAH1042 | *fyv6Δ* $Prp22^{WT}$ | Lipinski et al., 2023 |
| yAAH3558 | MATa ade2 cup1Δ::ura3 his3 leu2 lys2 trp1 ura3 GAL+ prp22Δ::loxP +pAAH1648 | $Prp22^{I1133R}$ | This study |
| yAAH3359 | MATa ade2 cup1Δ::ura3 his3 leu2 lys2 trp1 ura3 GAL+ prp22Δ::loxP fyv6Δ::hygR +pAAH1648 | *fyv6Δ* $Prp22^{I1133R}$ | This study |
| yAAH3606 | MATa ade2 cup1Δ::ura3 his3 leu2 lys2 trp1 ura3 GAL+ prp22Δ::loxP +pAAH1665 | $Prp22^{R805A}$ | This study |
| yAAH3607 | MATa ade2 cup1Δ::ura3 his3 leu2 lys2 trp1 ura3 GAL+ prp22Δ::loxP +pAAH1664 | $Prp22^{G810A}$ | This study |
| yAAH3608 | MATa ade2 cup1Δ::ura3 his3 leu2 lys2 trp1 ura3 GAL+ prp22Δ::loxP fyv6Δ::hygR +pAAH1664 | *fyv6Δ* $Prp22^{G810A}$ | This study |
| yAAH3612 | yAAH1930 + pAAH1675 | pre-5-FOA selection $Prp22^{R805A+I1133R}$ | This study |
| yAAH3613 | yAAH1930 + pAAH1674 | pre-5-FOA selection $Prp22^{G810A+I1133R}$ | This study |
| yAAH3614 | yAAH3377 + pAAH1665 | pre-5-FOA selection *fyv6Δ* $Prp22^{R805A}$ | This study |
| yAAH3615 | yAAH3377 + pAAH1675 | pre-5-FOA selection *fyv6Δ* $Prp22^{R805A+I1133R}$ | This study |
| yAAH3616 | yAAH3377 + pAAH1674 | pre-5-FOA selection *fyv6Δ* $Prp22^{G810A+I1133R}$ | This study |
| yAAH3632 | MATa ade2 cup1Δ::ura3 his3 leu2 lys2 trp1 ura3 GAL+ prp22Δ::loxP fyv6Δ::hygR +pAAH1675 | *fyv6Δ* $Prp22^{R805A+I1133R}$ | This study |
| yAAH3633 | MATa ade2 cup1Δ::ura3 his3 leu2 lys2 trp1 ura3 GAL+ prp22Δ::loxP fyv6Δ::hygR +pAAH1674 | *fyv6Δ* $Prp22^{G810A+I1133R}$ | This study |
| yAAH3635 | MATa ade2 cup1Δ::ura3 his3 leu2 lys2 trp1 ura3 GAL+ prp22Δ::loxP +pAAH1674 | $Prp22^{G810A+I1133R}$ | This study |
| yAAH3662 | MATa ade2 cup1Δ::ura3 his3 leu2 lys2 trp1 ura3 GAL+ prp22Δ::loxP +pAAH1675 | $Prp22^{R805A+I1133R}$ | This study |

| | | | |
|---|---|---|---|
| yAAH3663 | yAAH1931 + pAAH0470 | Prp22$^{WT}$ + WT ACT1-CUP1 | This study |
| yAAH3664 | yAAH1931 + pAAH0526 | Prp22$^{WT}$ + ACT1-CUP1 U301G | This study |
| yAAH3665 | yAAH1931 + pAAH0527 | Prp22$^{WT}$ + ACT1-CUP1 A302U | This study |
| yAAH3666 | yAAH3607 + pAAH0470 | Prp22$^{G810A}$ + WT ACT1-CUP1 | This study |
| yAAH3667 | yAAH3607 + pAAH0527 | Prp22$^{G810A}$ + ACT1-CUP1 A302U | This study |
| yAAH3694 | yAAH3607 + pAAH0526 | Prp22$^{G810A}$ + ACT1-CUP1 U301G | This study |
| yAAH3695 | yAAH3379 + pAAH0470 | *fyv6Δ* Prp22$^{WT}$ + WT ACT1-CUP1 | This study |
| yAAH3696 | yAAH3379 + pAAH0526 | *fyv6Δ* Prp22$^{WT}$ + ACT1-CUP1 U301G (gAG) | This study |
| yAAH3697 | yAAH3379 + pAAH0527 | *fyv6Δ* Prp22$^{WT}$ + ACT1-CUP1 A302U (UuG) | This study |
| yAAH3698 | yAAH3608 + pAAH0470 | *fyv6Δ* Prp22$^{WT}$ + WT ACT1-CUP1 | This study |
| yAAH3699 | yAAH3608 + pAAH0526 | *fyv6Δ* Prp22$^{WT}$ + ACT1-CUP1 U301G (gAG) | This study |
| yAAH3700 | yAAH3608 + pAAH0527 | *fyv6Δ* Prp22$^{WT}$ + ACT1-CUP1 A302U (UuG) | This study |
| yAAH3701 | yAAH1931 + pAAH0880 | Prp22$^{WT}$ + ACT1-CUP1 BSG | This study |
| yAAH3702 | yAAH3607 + pAAH0880 | Prp22$^{G810A}$ + ACT1-CUP1 BSG | This study |
| yAAH3703 | yAAH3379 + pAAH0880 | *fyv6Δ* Prp22$^{WT}$ + ACT1-CUP1 BSG | This study |
| yAAH3704 | yAAH3608 + pAAH0880 | *fyv6Δ* Prp22$^{G810A}$ + ACT1-CUP1 BSG | This study |
| yAAH3748 | yAAH3559 + pAAH0470 | *fyv6Δ* Prp22$^{I1133R}$ + WT ACT1-CUP1 | This study |
| yAAH3749 | yAAH3559 + pAAH0526 | *fyv6Δ* Prp22$^{I1133R}$ + ACT1-CUP1 U301G (gAG) | This study |
| yAAH3750 | yAAH3559 + pAAH0527 | *fyv6Δ* Prp22$^{I1133R}$ + ACT1-CUP1 A302U (UuG) | This study |
| yAAH3751 | yAAH3559 + pAAH0880 | *fyv6Δ* Prp22$^{I1133R}$ + ACT1-CUP1 BSG | This study |
| yAAH3771 | yAAH3635 + pAAH0470 | Prp22$^{G810A+I1133R}$ + WT ACT1-CUP1 | This study |
| yAAH3772 | yAAH3635 + pAAH0526 | Prp22$^{G810A+I1133R}$ + ACT1-CUP1 U301G | This study |
| yAAH3773 | yAAH3635 + pAAH0527 | Prp22$^{G810A+I1133R}$ + ACT1-CUP1 A302U | This study |
| yAAH3774 | yAAH3635 + pAAH0880 | Prp22$^{G810A+I1133R}$ + ACT1-CUP1 BSG | This study |
| yAAH3779 | yAAH3558 + pAAH0470 | Prp22$^{I1133R}$ + WT ACT1-CUP1 | This study |
| yAAH3780 | yAAH3558 + pAAH0526 | Prp22$^{I1133R}$ + ACT1-CUP1 U301G | This study |
| yAAH3781 | yAAH3558 + pAAH0527 | Prp22$^{I1133R}$ + ACT1-CUP1 A302U | This study |
| yAAH3782 | yAAH3558 + pAAH0880 | Prp22$^{I1133R}$ + ACT1-CUP1 BSG | This study |
| yAAH3783 | yAAH3633 + pAAH0470 | *fyv6Δ* Prp22$^{G810A+I1133R}$ + WT ACT1-CUP1 | This study |
| yAAH3784 | yAAH3633 + pAAH0526 | *fyv6Δ* Prp22$^{G810A+I1133R}$ + ACT1-CUP1 U301G | This study |

| | | | |
|---|---|---|---|
| yAAH3785 | yAAH3633 + pAAH0527 | *fyv6Δ* Prp22$^{G810A+I1133R}$ + ACT1-CUP1 A302U | This study |
| yAAH3786 | yAAH3633 + pAAH0880 | *fyv6Δ* Prp22$^{G810A+I1133R}$ + ACT1-CUP1 BSG | This study |
| BCY123 | MATa pep4::HIS3 prb1::LEU2 bar1::HIS6 lys2::GAL1/10-GAL4 can1 ade2 trp1 ura3 his3 leu2-3,112 | Protease-deficient yeast protein expression strain | Galej et al., 2013 |

**SupplementaryTable 7. Plasmids used in this study.**

| Plasmid ID | Plasmid name | Description | Source |
|---|---|---|---|
| pAAH0135 | pRS414 | CEN6/ARSH4 TRP1 Vector for Fyv6 plasmids | Mumberg et al., 1995 (ATCC# 87519) |
| pAAH1555 | pRS416-Fyv6 | FYV6 +/- ~250 bp (URA3 CEN6/ARSH4), used for Fyv6 shuffle strain | This study |
| pAAH1556 | pRS414-Fyv6 | FYV6 +/- ~250 bp (TRP1 CEN6/ARSH4) | This study |
| pAAH1572 | pRS414-FLAG-Fyv6 | N-terminally FLAG tagged FYV6 +/- ~250 bp (TRP1 CEN6/ARSH4) | This study |
| pAAH1577 | pRS414-FLAG-Fyv6-Δ1-16 | FLAG-Fyv6 Δ1-16 (TRP1 CEN ARS); in Fyv6 Δ1-16 truncation strain | This study |
| pAAH1578 | pRS414-FLAG-Fyv6-Δ1-23 | FLAG-Fyv6 Δ1-23 (TRP1 CEN ARS); in Fyv6 Δ1-23 truncation strain | This study |
| pAAH1579 | pRS414-FLAG-Fyv6-Δ1-51 | FLAG-Fyv6 Δ1-51 (TRP1 CEN ARS); in Fyv6 Δ1-51 truncation strain | This study |
| pAAH1580 | pRS414-FLAG-Fyv6-Δ134-173 | FLAG-Fyv6 Δ134-173 (TRP1 CEN ARS); in Fyv6 Δ134-173 truncation strain | This study |
| pAAH1581 | pRS414-FLAG-Fyv6-Δ103-173 | FLAG-Fyv6 Δ103-173 (TRP1 CEN ARS); in Fyv6 Δ103-173 truncation strain | This study |
| pAAH1573 | pRS414-Fyv6-FLAG | C-terminally FLAG tagged FYV6 +/- ~250 bp (TRP1 CEN6/ARSH4) | This study |
| pAAH1586 | pRS414-Fyv6-Δ103-173 -FLAG | Fyv6 Δ103-173-FLAG (TRP1 CEN ARS); in Fyv6 Δ103-173 truncation C-terminally FLAG tagged strain | This study |
| pAAH1602 | p360-Prp18 WT | $Prp18^{WT}$ (URA3 CEN) | Aronova et al., 2007; gift from Beate Schwer |
| pAAH1603 | p360-Prp18-11 | $Prp18^{V191A}$ (URA3 CEN) | Aronova et al., 2007; gift from Beate Schwer |
| pAAH1604 | p360-Prp18-18 | $Prp18^{S162P}$ (URA3 CEN) | Aronova et al., 2007; gift from Beate Schwer |
| pAAH1605 | p360-Prp18-11/18 | $Prp18^{S162P+V191A}$ (URA3 CEN) | Aronova et al., 2007; gift from Beate Schwer |
| pAAH0470 | ACT1-CUP1 WT | WT ACT1-CUP1 reporter; 38 nt BP-3' SS spacing. (GAP promoter, LEU2) | Gift from Charles Query. |
| pAAH1632 | ACT1-CUP1-9nt | ACT1-CUP1 reporter with 9 nt BP-3' SS spacing. (GAP promoter, LEU2) | This study |
| pAAH1633 | ACT1-CUP1-12nt | ACT1-CUP1 reporter with 12 nt BP-3' SS spacing. (GAP promoter, LEU2) | This study |
| pAAH1634 | ACT1-CUP1-21nt | ACT1-CUP1 reporter with 21 nt BP-3' SS spacing. (GAP promoter, LEU2) | This study |
| pAAH1635 | ACT1-CUP1-27nt | ACT1-CUP1 reporter with 27 nt BP-3' SS spacing. (GAP promoter, LEU2) | This study |
| pAAH1636 | ACT1-CUP1-15nt | ACT1-CUP1 reporter with 27 nt BP-3' SS spacing. (GAP promoter, LEU2) | This study |
| pAAH1637 | ACT1-CUP1-42nt | ACT1-CUP1 reporter with 42 nt BP-3' SS spacing. (GAP promoter, LEU2) | This study |
| pAAH1638 | ACT1-CUP1-46nt | ACT1-CUP1 reporter with 46 nt BP-3' SS spacing. (GAP promoter, LEU2) | This study |

| | | | |
|---|---|---|---|
| pAAH1639 | ACT1-CUP1-50nt | ACT1-CUP1 reporter with 50 nt BP-3' SS spacing. (GAP promoter, LEU2) | This study |
| pAAH1624 | pRS416-Syf1 | SYF1 +/- ~275 bp (URA3 CEN6/ARSH4), used for SYF1 shuffle strain | This study |
| pAAH1625 | pRS414-Syf1 | SYF1 +/- ~275 bp (TRP1 CEN6/ARSH4) | This study |
| pAAH1666 | pRS414-Syf1Δ817-859 | SYF1 Δ817-859 (TRP1 CEN6/ARSH4) | This study |
| pAAH1667 | pRS414-Syf1Δ778-859 | SYF1 Δ778-859 (TRP1 CEN6/ARSH4) | This study |
| pAAH1668 | pRS414-Syf1Δ634-859 | SYF1 Δ634-859 (TRP1 CEN6/ARSH4) | This study |
| pAAH1611 | pRS314-Cef1-WT | $Cef1^{WT}$ (TRP1 CEN) | Query and Konarska, 2012 ; Gift from Charles Query. |
| pAAH1612 | pRS314-Cef1-A37P | $Cef1^{A37P}$ (TRP1 CEN) | Query and Konarska, 2012; Gift from Charles Query. |
| pAAH1613 | pRS314-Cef1-S48R | $Cef1^{S48R}$ (TRP1 CEN) | Query and Konarska, 2012; Gift from Charles Query. |
| pAAH1614 | pRS314-Cef1-9-8 | $Cef1^{V36R}$ (TRP1 CEN) | Query and Konarska, 2012; Gift from Charles Query. |
| pAAH1641 | pRS314-Cef1-A37V | $Cef1^{A37V}$ (TRP1 CEN) | This study |
| pAAH1642 | pRS314-Cef1-M175I | $Cef1^{M175I}$ (TRP1 CEN) | This study |
| pAAH1643 | pRS314-Cef1-Q193P | $Cef1^{Q193P}$ (TRP1 CEN) | This study |
| pAAH1658 | PRS316-Cef1 | WT Cef1 (URA3 CEN) | Query and Konarska, 2012; Gift from Charles Query. |
| pAAH1440 | pRS424-Prp8 | WT Prp8 (full length) (TRP1 2μ) | This study |
| pAAH1659 | pRS424-Prp8-S1584Y | Prp8 S1584Y (TRP1 2μ) | This study |
| pAAH1660 | pRS424-Prp8-S1584F | Prp8 S1584F (TRP1 2μ) | This study |
| pAAH1661 | pRS424-Prp8-V1862L | Prp8 V1862L (TRP1 2μ) | This study |
| pAAH1662 | pRS424-Prp8-G1868R | Prp8 G1868R (TRP1 2μ) | This study |
| pAAH1663 | pRS424-Prp8-T1982S | Prp8 T1982S (TRP1 2μ) | This study |
| pAAH1042 | pPrp22-WT | WT Prp22 (TRP1 CEN) | Gift from Charles Query. |
| pAAH1648 | pPrp22-I1133R | Prp22 I1133R (TRP1 CEN) | This study |
| pAAH1664 | pPrp22-G810A | Prp22 G810A (TRP1 CEN) | Schwer and Meszaros, 2000; Gift from Beate Schwer |
| pAAH1665 | pPrp22-R805A | Prp22 R805A (TRP1 CEN) | Schwer and Meszaros, 2000; Gift from Beate Schwer |

| | | | |
|---|---|---|---|
| pAAH1674 | pPrp22-G810A+I1133R | Prp22 G810A+I1133R (TRP1 CEN) | This study |
| pAAH1675 | pPrp22-R805A+I1133R | Prp22 R805A+I1133R (TRP1 CEN) | This study |
| pRS424-CBP-His-TEV-Prp22-S635A | pRS424-CBP-His-TEV-Prp22-S635A | GAL/GAPDH promoter, Prp22 expression (TRP1 2µ) | This study |
| pRS426-CBP-His-TEV-Prp22-S635A | pRS426-CBP-His-TEV-Prp22-S635A | GAL/GAPDH promoter, Prp22 expression (URA3 2µ) | This study |
| pAAH0880 | ACT1-CUP1 BSG (A259G) | BS A259G reporter used for ACT1-CUP1 assays. (GAP promoter, LEU2) | Gift from Charles Query. |
| pAAH0526 | ACT1-CUP1 U301G | 3' SS gAG reporter used for ACT1-CUP1 assays. (GAP promoter, LEU2) | Gift from Charles Query. |
| pAAH0527 | ACT1-CUP1 A302U | 3' SS UuG reporter used for ACT1-CUP1 assays. (GAP promoter, LEU2) | Gift from Charles Query. |

**Supplementary table 8. Oligonucleotides used in this study.**

| Designation | Sequence (5′ to 3′) | Use | Source or reference |
|---|---|---|---|
| SUS1-exon1 | TGGATACTGCGCA ATTAAAGAGTC | RT-PCR primer | Jacewicz et al., 2015; doi: 10.1261/rna.048942 .114 |
| SUS1-exon3 | TCATTGTGTATCTA CAATCTCTTCAAG | RT-PCR primer | Jacewicz et al., 2015; doi: 10.1261/rna.048942 .114 |
| YOS1-F | AGT ACT GAG CGA AGA AAG | RT-PCR primer | This paper |
| YOS1-R | CAT CCC AAT AGT AAT TCA TAA AC | RT-PCR primer | This paper |
| RPS18A_fwd | ACAAGGTTCCTTCC AACACA | RT-PCR primer | This paper |
| RPS18A_rev | TAACGACGACCAAC ACCCTT | RT-PCR primer | This paper |
| CGI121_fwd | GACAAAGAGCAATT GAGGACGA | RT-PCR primer | This paper |
| CGI121_rev | CGTCTTGGGGCTTG AAATCA | RT-PCR primer | This paper |

| | | | |
|---|---|---|---|
| NMD2_fwd | CTGCATTGATAATA CATTGGACAGA | RT-PCR primer | This paper |
| NMD2_rev | GAACCTTTCACAAA ACCCTTCT | RT-PCR primer | This paper |
| OST5_fwd | CAAGGTTTACAATG ACTTATGAACA | RT-PCR primer | This paper |
| OST5_rev | ACAGGGCAGATGAT AATACAGC | RT-PCR primer | This paper |
| DID4_fwd | GGAAAGAATGTCAC TCCGCA | RT-PCR primer | This paper |
| DID4_rev | CTTGCCCATTCTTT GCCGAT | RT-PCR primer | This paper |
| RPS7B_fwd | TCGCCAAGACCTTC ATCGAT | RT-PCR primer | This paper |
| RPS7B_rev | AGACAAAGCTGGAA CTGGGA | RT-PCR primer | This paper |
| yU6 | /5IRD700/GAACTGC TGATCATCTCTG | For primer extension | Xu and Query, 2007; doi: 10.1016/j.molcel.20 07.09.022 |
| YAC6 | /5IRD700/GGCACTC ATGACCTTC | For primer extension | Siatecka et al., 1999; doi: 10.1101/gad.13.15.1 983 |

| | | | |
|---|---|---|---|
| UBC4-complementary oligo | ATGAAGTAGGTGGAT | for RNase H cleavage | Wilkinson et al., 2017; DOI: 10.1126/science.aar3729 |

**SUPPLEMENTARY METHODS**

The following flags were used with the specified tools for RNA-seq data analysis.

Assessment of RNA-seq quality: FASTQC

Trimming: fastp --detect_adapter_for_pe -Q -q 20 -u 40 -l 36 --poly_g_min_len 10 -g -5 -3 -W 4 -M 20

Saccer3 Annotation: Ensembl, R64-1-1

Read Mapping: All samples were aligned with STAR in a first pass. Novel junctions from all samples (WT and *fyv6Δ*) were combined and filtered for likely false-positive junctions (non-canonical junctions, column5 > 0; junctions supported by multi mappers only, column 7 > 0; junctions supported by two few reads, column 7 > 2). STAR was run in a second pass with the additional input of the filtered set of novel junctions as follows: --runMode alignReads --runThreadN 20 --genomeDir ./genome --sjdbGTFfile annotation.gtf --alignIntronMin 10 --alignIntronMax 2000 --readFilesCommand zcat --readFilesIn "R1_trimmed.fastq.gz" "R2_trimmed.fastq.gz" --outSAMtype BAM SortedByCoordinate --limitBAMsortRAM 6000000000

SpliceWiz Differential Splicing Analysis: with the following flags: rmats.py -t paired --readLength 150 --variable-read-length --novelSS.

FeatureCounts Split Read Counting: featureCounts -a annotation.gtf -o Sample --splitOnly -J -f -p -T 5 -R BAM -B -C Aligned.sortedByCoord.out.bam

Salmon Read Count Quantitation: Salmon was run on the free resource usegalaxy.org with the following flags: --libType A --incompatPrior '0.0' --biasSpeedSamp '5' --fldMax '1000' --fldMean '250' --fldSD '25' --forgettingFactor '0.65' --maxReadOcc '100' --numBiasSamples '2000000' --numAuxModelSamples '5000000' --numPreAuxModelSamples '5000' --numGibbsSamples '0' --numBootstraps '0' --thinningFactor '16' --sigDigits '3' --vbPrior '1e-05'. Differential gene expression analysis was conducted with DeSeq2with the following flags: -t 1 -P -V 10 -i -y salmon -x mapping.gff.

DeSeq2 Differential Gene Expression Analysis: deseq2.R -o 'output.dat' -p -A 0.1 -H -f '[["Fyv6", [{"WT": ["WT1.tabular", "WT2.tabular"]}, {"Fyv6": ["Fyv61.tabular", "Fyv62.tabular"]}]]]' -l '{"Fyv61.tabular": "Fyv61.tabular", "Fyv62.tabular": "Fyv62.tabular", "WT1.tabular": "WT1.tabular", "WT2.tabular": "WT2.tabular"}' -t 1 -P -V 10 -i -y salmon -x

FAnS Calculation: Junction reads within genes were counted with featureCounts. Genes were filtered based on those listed in the Ares Intron database. Canonical 5′ and 3′ SS were annotated based on the highest number of junction counts. All junctions were filtered based on presence in all four RNA-seq samples. Read counts for filtered junctions were combined for *fyv6Δ* and WT replicates. Alternative junctions were separated into those that shared a canonical 5′ SS and an alternative 3′ SS or those that shared a canonical 3′ SS and an alternative 5′ SS. FAnS was calculated based on the number of junction reads for a unique alternative 3′ SS sharing a canonical 5′ SS divided by the number of junction reads for canonical 5′ SS and 3′ SS within a sample to adjust for changes in expression. Ratios of FAnS were calculated by dividing the FAnS value for *fyv6Δ* by the FAnS value for WT.

Docker Images

fastp - biocontainers/fastp

STAR - alexdobin/star:2.7.10a_alpha_220506

featureCounts - pegi3s/feature-counts:2.0.0

Salmon - combinelab/salmon:1.10.3

FASTQC - jysgro/fastqc:ub2306_12.1
